# AGC kinase members phosphorylate ubiquitin

**DOI:** 10.1101/2020.07.15.204149

**Authors:** Carl-Christian Kolbe, Eicke Latz

## Abstract

The posttranslational modification of proteins with ubiquitin controls most cellular processes, such as protein degradation or transport, cell signaling, or transcription^1–5^. Ubiquitin can be phosphorylated at multiple sites, which likely further modulates the function of protein ubiquitination^6,7^. However, except for PINK1^8–10^, the kinases involved in ubiquitin phosphorylation remain unknown, which hampers our understanding of phospho-ubiquitin signaling. In this study, we performed genome-wide in vitro kinase screenings and discovered that AGC kinases phosphorylate ubiquitin. Ubiquitin phosphorylation by members of the PKA, PKC, PKG and RSK families as well as by less well-characterized kinases, such as SGK2, was not solely dependent on peptide specificity but required additional kinase recruitment to ubiquitin. The stabilization of the kinase interaction with ubiquitin resulted in phosphorylation of suboptimal kinase motifs on ubiquitin, suggesting that ubiquitin phosphorylation is dictated primarily through the recruitment of kinases to the ubiquitinated proteins. Hence, we identify AGC kinase members as enzymes that can phosphorylate ubiquitin in a mechanism regulated by protein interactions outside of the catalytic kinase domain and are only applicable to specific subsets of ubiquitinated proteins.

## Main

Ubiquitin phosphorylation represents an expansion of the classical ubiquitin code in which the timely addition and removal of phosphoryl groups by kinases and phosphatases modulates the function of protein ubiquitination^6,7^. Proteomic analysis showed that ubiquitin gets phosphorylated at T7, T12, T14, S20, T22, S57, Y59, S65, and T66 (Extended Data Fig. 1a). To understand the regulation of phospho-ubiquitination and elucidate its biological function, it is crucial to identify the enzymes that regulate ubiquitin modifications. Indeed, the identification of PINK1-dependent phosphorylation of ubiquitin at S65^8–10^ was a breakthrough discovery, which clarified how phospho-ubiquitination regulates physiological and pathophysiological processes^11–15^. However, which other kinases facilitate ubiquitin phosphorylation remain unknown^6,7,16^. Conceivably, phosphorylation of other amino acids within ubiquitin will significantly modify the ubiquitin surface and thereby modulate ubiquitin function in similar ways as S65 phosphorylation^6,7,16^. Therefore, we performed an unbiased in vitro kinase screening to identify the kinases that can phosphorylate ubiquitin.

**Figure 1:**
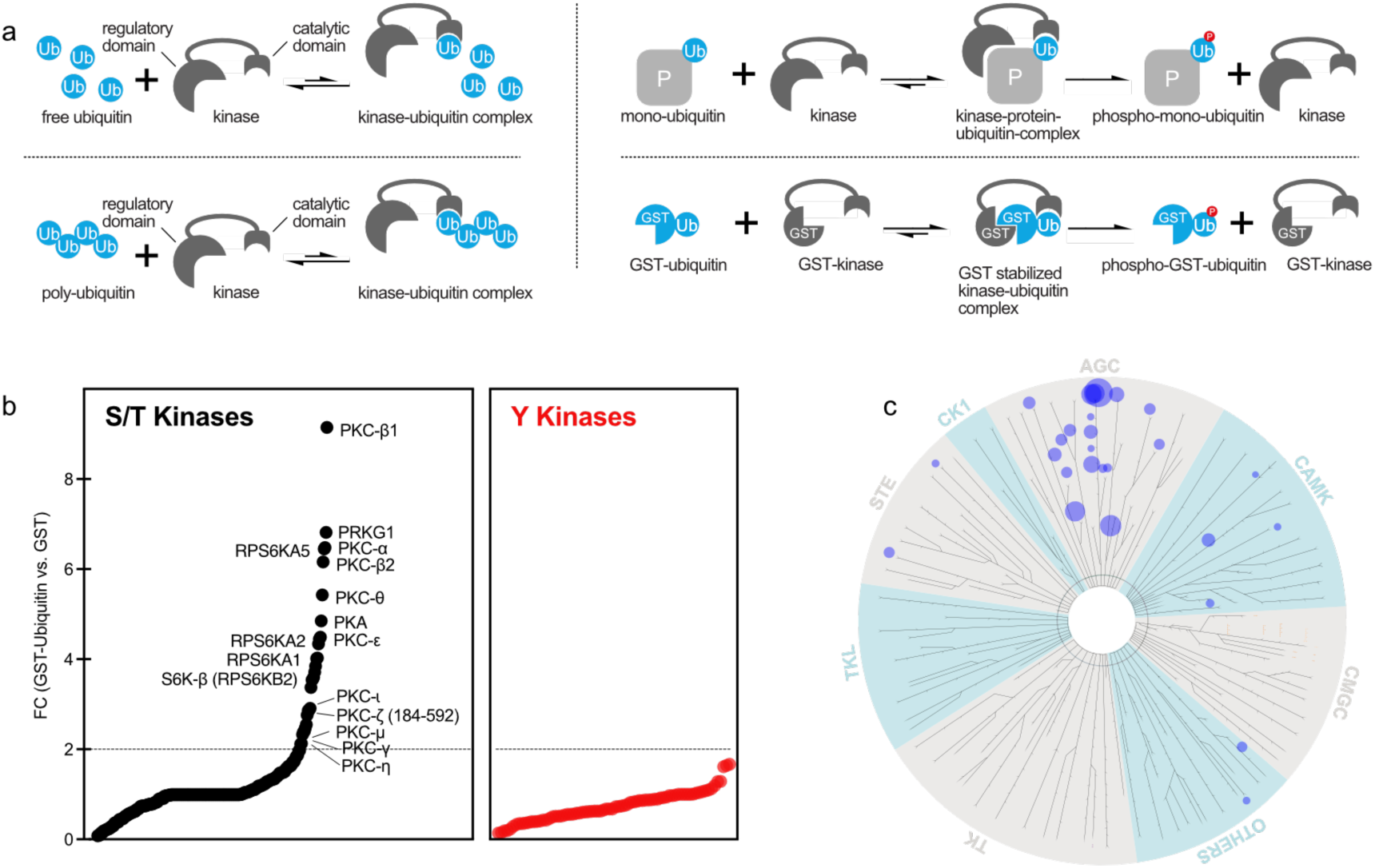
GST-Ubiquitin is phosphorylated by members of the AGC kinase group. (a) Schematic representation of kinase recruitment to free-, poly- and substrate-bound ubiquitin. Ubiquitin phosphorylation is only achieved if the regulatory kinase domain can bind to the ubiquitinated protein (P) to create a stable ubiquitin-kinase complex. GST based dimerization between kinase and ubiquitin is displayed in last panel. (b) [^33^P] incorporation indicated as fold change (FC) between GST-ubiquitin and GST as control. S/T kinases are shown in black. Y kinases are shown in red. Selected candidate kinases are named individually. Kinase screening was performed once. (c) Kinases with FC > 2, between GST-ubiquitin and GST, are mapped on a phylogenetic kinase tree as blue dots. Dot size refers to FC measured in kinase screening.

Two essential parameters control protein phosphorylation, i.e., peptide specificity and kinase recruitment. Peptide specificity is a kinase’s ability to phosphorylate a specific linear sequence of amino acids, whereas kinase recruitment describes the interaction between substrate and regulatory kinase domains (Extended Data Fig. 1b). Notably, strong kinase recruitment to the substrate can overcome poor peptide specificity^17^. Hence, protein-protein interaction outside of the catalytic kinase domain represents a crucial parameter of protein phosphorylation. In contrast to most other proteins, ubiquitin is present in three distinct forms inside the cell, i.e., free ubiquitin, polyubiquitin and protein-ubiquitin conjugats^18^, which all might affect their ability to serve as kinase substrates (Fig. 1a). Since the conjugation of ubiquitin to proteins could facilitate the interaction between a kinase and ubiquitin, we hypothesized that kinase recruitment could be of particular importance for ubiquitin phosphorylation. Such a ubiquitin substrate-dependent phosphorylation could significantly alter kinase efficiency depending on whether free-, poly- or protein-bound ubiquitin is available. Subsequently, kinase recruitment facilitated by the ubiquitinated protein would result in highly specific ubiquitin phosphorylation. Such precise regulation of ubiquitin phosphorylation could modify specific protein-ubiquitin conjugates while leaving the majority of ubiquitin unaltered.

### Ubiquitin phosphorylation by AGC kinases

To identify kinases responsible for ubiquitin phosphorylation, we performed a genome-wide in vitro kinase screening, including 339 recombinant kinases, of which 278 were N-terminally tagged with GST (Extended Table 1). After incubating recombinant monomeric ubiquitin and the individual kinases with [^33^P]-γ-ATP using optimized kinase reaction parameters for each kinase, we measured [^33^P] incorporation to assess kinase activity. To test for the effect of increased kinase-substrate interaction on ubiquitin phosphorylation, we performed the kinase screening assays with both unmodified ubiquitin and GST-tagged ubiquitin (Extended Data Fig. 1c). GST is an affinity tag, which leads to dimerization and creates an interaction between GST- tagged kinases and GST-ubiquitin^19^. GST-dimerization thereby mimics the interactions between ubiquitinated proteins and a regulatory kinase domain (Fig. 1a).

Kinase screening using untagged ubiquitin as a substrate did not reveal kinases that we could validate to phosphorylate ubiquitin (Extended Data Fig. 1d,e,f-j). In contrast we identified several kinases that phosphorylated GST-ubiquitin but not the GST control protein (Fig. 1b, Extended Data Fig. 2). Phylogenetic mapping of the identified GST-ubiquitin kinases showed that these kinases primarily clustered in the AGC-kinase group, including PKA (PRKACA), PKG (PRKG1), PKCs (PKC-α; PKC-β1; PKC-β2; PKC-θ, PKC-ε) and the structurally related MSKs (RPS6KA5) and RSKs (RPS6KA1; RPS6KA2) (Fig. 1b,c). Some members of the AGC kinase group are among the best-investigated protein kinases and are involved in numerous essential cellular signaling pathways. Second messengers, such as cyclic nucleotides (cAMP, cGMP), ions (Ca^2+^) or lipids (diacylglycerol) are known to regulate these kinases, and there is a high conservation in these mechanisms even found in early eukaryotic cells^20,21^.

**Figure 2:**
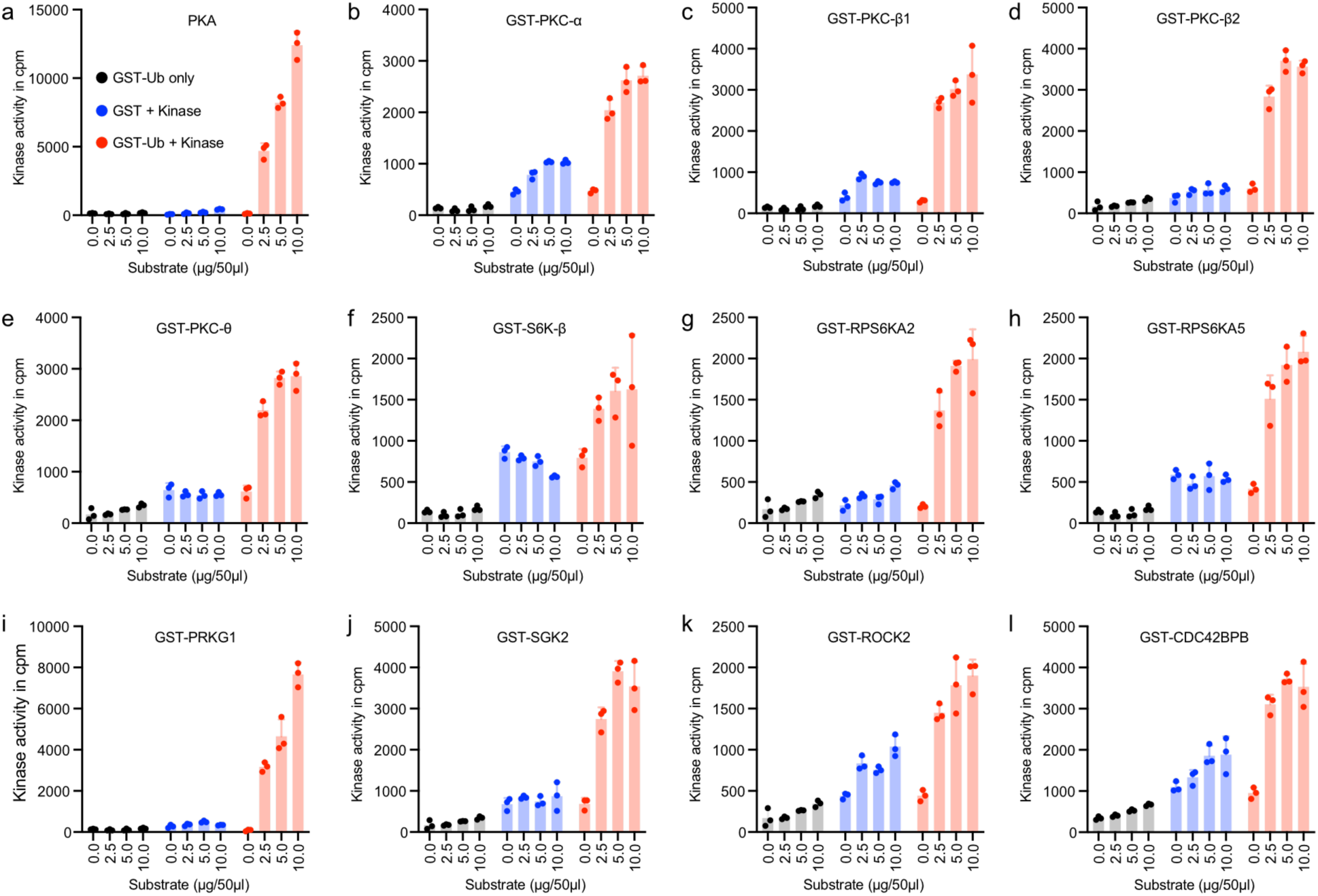
GST-ubiquitin is phosphorylated in concentration dependent manner. (a-l) Candidate kinases were tested with increasing amounts of GST-ubiquitin or GST (0 – 10 µg/ 50 µL). Shown is the [^33^P] incorporation in counts per minute (cpm). GST-ubiquitin without kinase is shown in grey, GST plus kinase in blue and conditions with GST-ubiquitin plus kinase in red. (individual data points are shown, bar graph: mean + SD, n= 3)

Several of these candidate AGC kinases were subsequently validated by GST and GST-ubiquitin titration to confirm substrate phosphorylation (Fig. 2a-l). In total, at least twelve AGC kinases showed ubiquitin-specific activity, as demonstrated by elevated [^33^P] incorporation for GST- ubiquitin compared to the GST control. Of all validated kinases only MAPKAP3, which had an FC > 4 in the screening, was identified as a GST instead of GST-ubiquitin specific kinase (Extended Data Fig. 3a, b). Furthermore, a selection of oncogenic kinases demonstrated no ubiquitin phosphorylation (Extended Data Fig. 4a-d).

**Figure 3:**
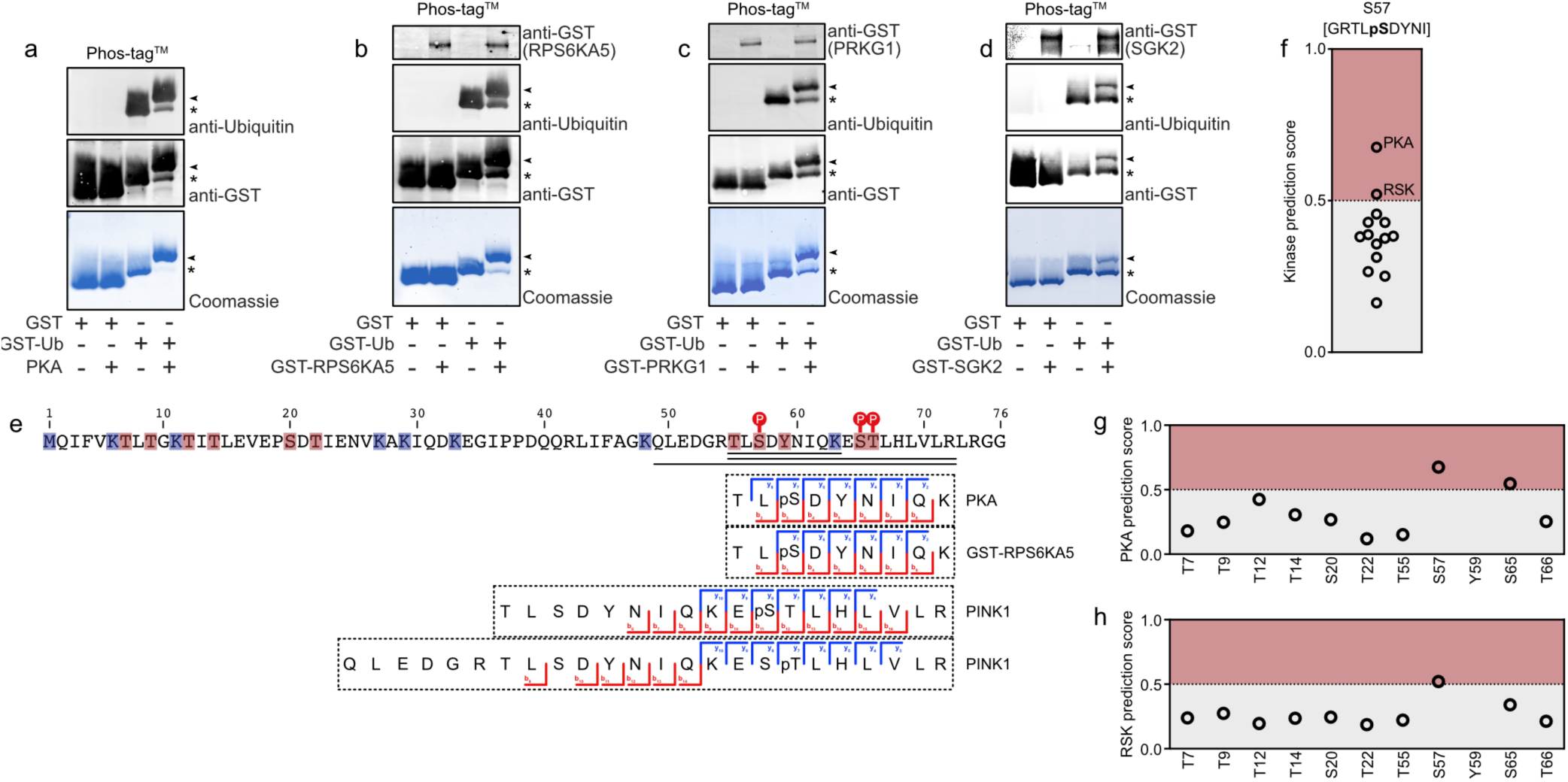
AGC kinase group members phosphorylate ubiquitin at S57. (a-d) Western blot and Coomassie analysis using Phos-tag^™^ SDS-PAGE to separate phosphorylated GST-ubiquitin (GST-Ub) after incubation with PKA, GST-PRKG1, GST-RPS6KA5 or GST-SGK2. Arrows indicates phosphorylated GST-ubiquitin; Asterisks indicates non- phosphorylated GST-ubiquitin. Shown are representative gels and Immunoblots from at least three independent experiments. (e) Identification of phosphorylated ubiquitin peptides by mass spectrometry. B- and y-ions upon treatment with indicated kinases are indicated. Phosphorylated amino acids are represented as pS or pT. (f) In silico S/T kinase prediction for S57 using NetPhos 3.1 server. Kinases with a prediction score > 0.5 are highlighted in red. (g,h) PKA and RSK in silico phosphorylation prediction for each S/T within ubiquitin using NetPhos 3.1 server. Phosphorylation sites with a prediction score > 0.5 are highlighted in red.

**Figure 4:**
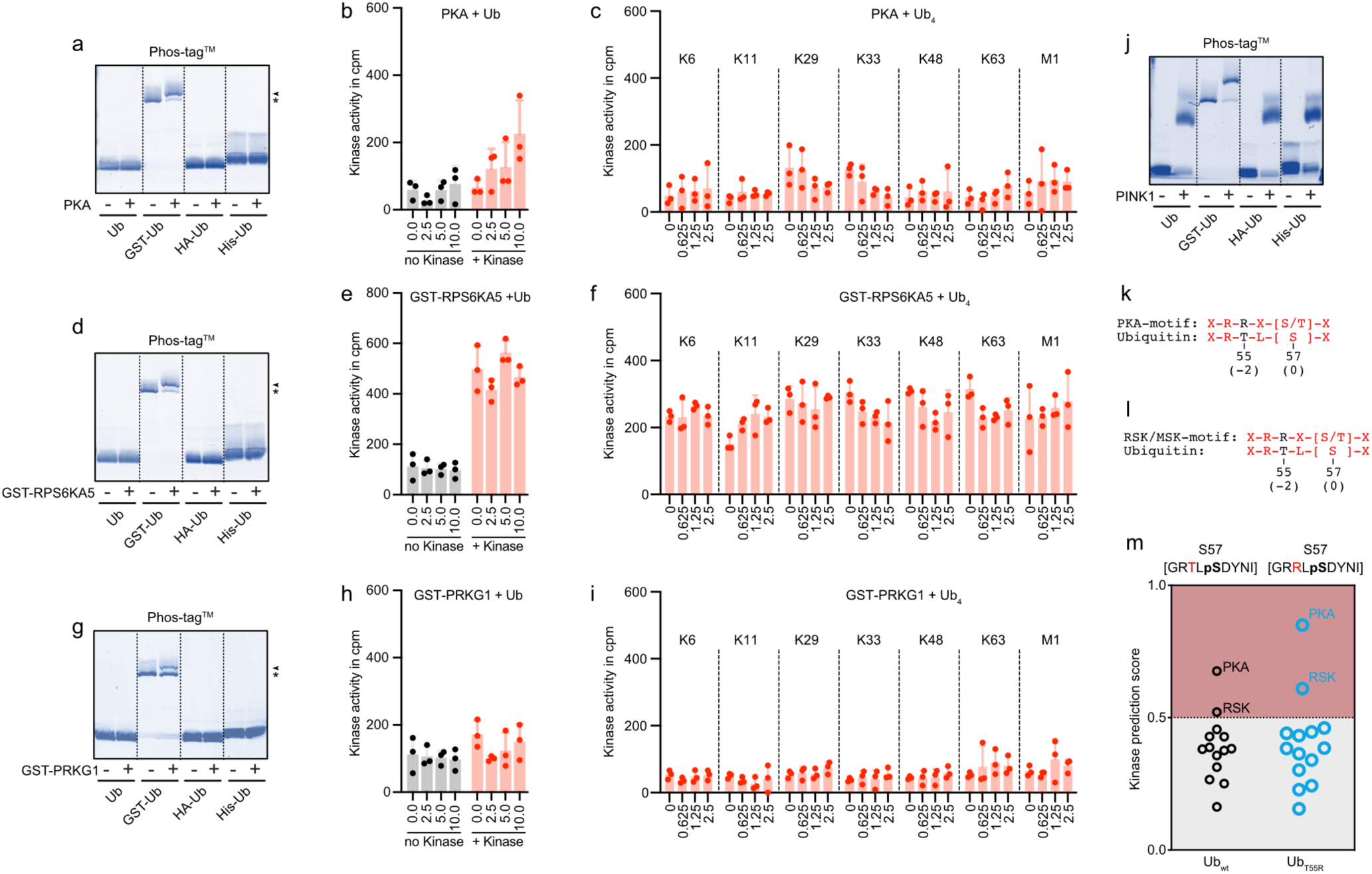
Kinase recruitment is essential for ubiquitin phosphorylation. (a,d,g) Phosphorylation of untagged, GST-, HA- or His-tagged ubiquitin assessed by Phos-tag^™^ SDS-PAGE. Phos-tag^™^ SDS-PAGE was used to separate phosphorylated ubiquitin proteins after incubation with PKA, GST-PRKG1 and GST-RPS6KA5. Arrow indicated phosphorylated GST-ubiquitin; Asterisk indicates non-phosphorylated GST-ubiquitin. Shown are representative gels from at least three independent experiments. (b,e,h) PKA, GST-PRKG1 and GST-RPS6KA5 were tested with increasing amounts of untagged ubiquitin (0 - 10 µg/ 50 µL). Shown is the [^33^P] incorporation in cpm. Ubiquitin without kinase is shown in grey and ubiquitin plus indicated kinase in red. (individual data points are shown, bar graph: mean + SD, n= 3). (c,f,i) Linkage type-specific tetra-ubiquitin (Ub_4_) phosphorylation by PKA, GST-PRKG1 and GST-RPS6KA5 were tested with increasing amounts of Ub_4_ (0 - 2.5 µg/ 50 µL). Y-axis displays [^33^P] incorporation in cpm. (individual data points are shown, bar graph: mean + SD, n= 3). (j) Phosphorylation of untagged, GST-, HA- or His-tagged ubiquitin by PINK1 was analyzed by Phos-tag^™^ SDS-PAGE. Shown is a representative gel from four independent experiments. (k,l) Alignment of PKA and RSK phosphorylation motifs with the amino acid sequence surrounding S57 of ubiquitin. (m) In silico S/T kinase prediction for S57 using NetPhos 3.1 server for wild type Ubiquitin T55R(Ub_T55R_). PKA and RSK have a phosphorylation prediction score >2 for Ub_wt_ and Ub_T55R_.

To quantify the amount of phosphorylated GST-ubiquitin, we analyzed kinase reactions on Phos-tag^™^ acrylamide SDS-PAGE to separate phosphorylated and non-phosphorylated ubiquitin^22^ (Extended Data Fig. 5). We identified a clear size shift for GST-ubiquitin after treatment with either PKA, GST-RPS6KA5, GST-PRKG1, or GST-SGK2, suggesting specific phosphorylation of the substrate. Furthermore, incubation of GST-ubiquitin with PKA, GST- RPS6KA5, GST-PRKG1 resulted in more than 50% of GST-ubiquitin phosphorylation by the respective kinases (Fig. 3a-d).

We performed fingerprint mass-spectrometry to identify the phosphorylation site on ubiquitin for the tested kinases and located the phosphorylation site attached by PKA and GST-RPS6KA5 to S57. In accordance with previous research, PINK1 exclusively phosphorylated S65^8–10^ and T66^23^ (Fig. 3e, Extended Data Fig. 6a-d). We did not observe phosphorylation of the control GST-tag by PKA, GST-RPS6KA5, or PINK1 (data not shown). S57 phosphorylation of ubiquitin is the most abundant ubiquitin phosphorylation under steady-state conditions, it was the first discovered ubiquitin phosphorylation site^24^, and large scale proteomic approaches identified this phosphorylation site more often than any other ubiquitin phosphorylation sites (Extended Data Fig. 6e). Of note, NetPhos^25,26^ in silico prediction supported our findings by ranking PKA and RSK families as the most likely kinases to phosphorylate S57 (Fig. 3f) and identified S57 as the preferred phosphorylation site for PKA and RSKs within ubiquitin (Fig. 3g,h). Together, these data validate that the respective members of the AGC-kinase group phosphorylate ubiquitin.

### Kinase recruitment is essential for ubiquitin phosphorylation

The kinase screening using untagged ubiquitin identified no ubiquitin targeting kinases. Likewise, the same kinases, which showed robust phosphorylation of GST-ubiquitin, did not phosphorylate untagged ubiquitin (Extended Figure 7a). We hypothesized that ubiquitin phosphorylation requires kinase recruitment. Therefore, we performed kinase reactions with untagged, GST-, HA- or His-tagged ubiquitin. As shown by Phos-tag^™^ SDS-PAGE, PKA, GST- RPS6KA5, and GST-PRKG1 phosphorylated only GST-ubiquitin. Due to the lack of kinase recruitment, no phosphorylation was observed for untagged ubiquitin and ubiquitin with non- dimerizing protein-tags, such as HA- or His-ubiquitin (Fig. 4a,d,g). Highly sensitive [^33^P] incorporation measurements failed to demonstrate phosphorylation of untagged ubiquitin (Fig. 4b,e,h). In accordance with the data for untagged ubiquitin, also tetra ubiquitin chains (Ub_4_) with K6, K11, K29, K33, K48, K63, and M1 specific linkage types (Extended Data Fig. 7b,c) were not phosphorylated by any of the indicated kinases due to the lack of kinase recruitment (Fig. 3c,f,i). In contrast, ubiquitin phosphorylation by PINK1 was independent of the N- terminal tag (Fig. 4j). To exclude that GST-induced ubiquitin phosphorylation was a result of allosteric kinase activation, GST, untagged ubiquitin, and GST-ubiquitin were pooled and incubated with either PKA or GST-RPS6KA5. Again, we detected GST-ubiquitin phosphorylation, whereas neither GST or ubiquitin nor the combination of both showed any signs of phosphorylation (Extended Data Fig. 7d).

The need for kinase recruitment for ubiquitin phosphorylation suggests that the phosphorylation motif surrounding S57 demonstrates relatively poor peptide specificity for the identified kinases. The classical phosphorylation motifs for PKA and RSKs/MSKs display two positively charged amino acids at position [-2] and [-3] upstream of the phosphorylated serine or threonine. Aligning the amino acid sequence around S57 with the classical peptide recognition motif for PKA and RSKs/MSKs shows that ubiquitin contains only a single arginine at position ([-3]/ R54) and is lacking a second positive amino acid at position ([-2]/ T55) (Fig. 4k,l). This suboptimal kinase motif appears insufficient to induce phosphorylation solely by peptide specificity but requires additional stabilization of the kinase ubiquitin complex by the ubiquitinated protein. Exchanging T55 to arginine (T55R) increased the likelihood of S57 to be phosphorylated by PKA and RSKs (Fig. 4m), while not affecting the phosphorylation prediction for any other phosphorylatable sites within ubiquitin (Extended Data Fig. 8a,b). These data strongly suggest that ubiquitin phosphorylation by a number of kinases requires strong enzyme-substrate interaction to overcome suboptimal kinase motifs.

## Discussion

A great variety of posttranslational modifications regulate protein function, protein longevity and protein-protein interactions. The addition or removal of protein modifications in time and space is dictated by the activity of signaling pathways. The identification which enzymes control posttranslational protein modifications has provided key information to understand cellular processes and their connection to the activity of a given signaling pathway. The fact that the protein posttranslational modification ubiquitin can be further posttranslationally modified itself by ubiquitin, ubiquitin-like proteins, phosphorylation or other modifications shows the complexity with which the so-called ubiquitin code affects eukaryotic biology. Ubiquitin is abundantly present in all eukaryotic cells and given the critical role of ubiquitination in cellular function it is expected that ubiquitin modification itself needs to be tightly controlled to prevent accidental addition of ubiquitin posttranslational modification. Indeed, in this study we showed, that a panel of 339 recombinant human kinases cannot phosphorylate free ubiquitin. However, we discovered that protein-conjugated ubiquitin makes a *bona fide* substrate motif for several kinases that integrate the activity of many classes of signaling receptors, such as GPCRs or innate immune receptor signaling families. We identified twelve kinases which belong to the AGC kinase group to phosphorylate ubiquitin. In contrast to PINK1, the thus far only known ubiquitin kinase, the members of the AGC kinase group require ubiquitin to be conjugated at a protein, which recruits the kinase to ubiquitin. Due to kinase recruitment, the suboptimal kinase motifs on ubiquitin can be overcome and ubiquitin can be phosphorylated. The prerequisite of kinase recruitment to ubiquitinated proteins for ubiquitin phosphorylation suggests that the sequential activities of ubiquitin ligases and ubiquitin kinases prevents this risk of accidental ubiquitin phosphorylation and allows for highly coordinated phosphorylation of ubiquitin. Curiously, the - as of yet only identified ubiquitin kinase - PINK1, can phosphorylate ubiquitin even in its free form. However, PINK1 activity itself is regulated by mitochondrial membrane depolarization and upon induction it is recruited to the outer mitochondrial membrane and is thereby spatially limited to substrates at the mitochondria surface^11^.

Including the ubiquitinated protein as a requirement for the process of ubiquitin phosphorylation allows for pinpointed ubiquitin phosphorylation driven by other pathways in which kinases are for example regulated by second messengers. Indeed, many signaling pathways that integrate mainly cell extrinsic signals, such as a GPCRs, provoke transient production of signaling molecules^27^, which is in line with the notion that the overall stoichiometry of S57 phosphorylated ubiquitin in resting cells is rather small^28^. The universal expression of AGC kinase members (Extended Data Fig. 9a,b), as well as countless publications focusing on these kinases, clearly demonstrate the significant role of these enzymes in nearly all biological processes. Due to the omnipresence of AGC kinases and ubiquitin, it is conceivable that S57 ubiquitin phosphorylation affects numerous signaling events but especially in situations where AGC kinases get activated by increased levels of their respective second messengers (e.g. cAMP, cGMP, or Ca^2+^).

The presence of at least twelve ubiquitin-specific kinases indicates a high evolutionary effort to precisely control ubiquitin phosphorylation at S57, demonstrating its importance for correct cellular function. Identifying AGC kinase members as ubiquitin-specific kinases is completing our understanding of how the ubiquitin code is written and creates innumerous possibilities for future research. Pharmaceutical research could especially benefit from our results to manipulate the ubiquitin proteasome system (UPS). Although the UPS is a highly promising field for future drug development, the overall number of FDA-approved UPS drugs still lacks behind those of kinase inhibitors^29,30^. Combining the already existing kinase inhibitors with newly developed UPS drugs could lead to promising combinatorial therapies in which both drug classes are used synergistically. However, even more important is the observation that ubiquitin phosphorylation depends heavily on kinase recruitment by the ubiquitinated protein, which adds a novel and unexpected level of complexity to the ubiquitin code. Future research should focus on identifying ubiquitin-specific kinases for individual ubiquitinated proteins.

## Acknowledgments

We thank T. Ruppert, S. Merker and U. Bach (Core Facility for Mass Spectrometry & Proteomics, Zentrum für Molekulare Biologie der Universität Heidelberg) for their assistance with mass spectrometry. We thank F. Totzke (ProQinase GmbH) for his implementation and execution of radioactive in vitro kinase assays. Further we would like to thank M. Geyer (Institute of Structural Biology, University of Bonn) for his constructive criticism of the manuscript. The work was funded by the Deutsche Forschungsgemeinschaft (DFG, German Research Foundation) under Germany’s Excellence Strategy – EXC2151 – 390873048”. E. Latz is co-founder of IFM Therapeutics. The authors declare no additional competing financial interests.

## Author contributions

C.C. Kolbe and E. Latz designed the study and wrote the manuscript. C.C. Kolbe performed experiments and analyzed data.

## Methods

### Kinases

A detailed list of kinases, including assay concentrations used in this study for radioactive in vitro kinase assays can be obtained from Supplementary Data Table 1.

### Protein Preparation

Glutathione S-Transferase (GST) (GenScript; Cat. # Z02039-1), ubiquitin (R&D Systems; Cat. # U-100H), His-ubiquitin (R&D Systems; Cat. # U-530), HA-ubiquitin (R&D Systems; Cat. # U-110) or GST-ubiquitin (R&D Systems; Cat. # U-540) were solubilized in 50 mM HEPES (pH= 7.4), if not otherwise stated. Buffer exchange was performed using Zeba™ spin desalting columns, 7K MWCO (Thermo Fisher; Cat. # 89883). AQUA pure tetra ubiquitin chains were purchased from R&D Systems (Cat. # UC-15; UC-45; UC-83; UC-103; UC-210B; UC-310B; UC-710B; UC-750), and purity was confirmed by SDS-PAGE.

### Radioactive in vitro kinase assay

Standard kinase reaction conditions: 70 mM HEPES-NaOH with pH= 7.5 (Sigma; Cat. # H3375), 3 mM MgCl_2_ (Sigma; Cat. # M3634), 3 mM MnCl_2_ (VWR; Cat. # 1.05927.1000), 3 µM Na- orthovanadate (Sigma; Cat. # S6508), 1.2 mM DTT (Sigma; Cat. # D-0632), 50 µg/mL PEG_20,000_ (SERVA; Cat. # 33138), ATP (Sigma; Cat. # A-7699) (variable, corresponding to ATP-K_m_ of the respective kinase, [^33^P]- γ-ATP (PerkinElmer; Cat. # NEG302H): approx. 7-8 × 10^5^ cpm, 1 % (v/v) DMSO (Sigma; Cat. # 154938), recombinant protein kinase (variable, indicated in Extended Data Table 1). All PKC assays (except the PKC-mu and the PKC-nu assay) additionally contained 1 mM CaCl_2_ (VWR; Cat. # 1.02382), 4 mM EDTA (Roth; Cat. # 8040.1), 5 μg/ml Phosphatidylserine (Fluka; Cat. # 79405) and 1 μg/ml 1,2-Dioleylglycerol (Sigma; Cat. # D8394). The CAMK1D, CAMK2A, CAMK2B, CAMK2D, CAMK4, CAMKK1, CAMKK2, DAPK2, EEF2K, MYLK, MYLK2 and MYLK3 assays additionally contained 1 μg/ml Calmodulin (Millipore; Cat. # 14-368) and 0.5 mM CaCl_2_ (VWR, Cat. # 1.02382). The PRKG1 and PRKG2 assays additionally contained 1 μM cGMP (Sigma; Cat. # G-6129). The DNA-PK assay additionally contained 2.5 μg/ml DNA (Sigma; Cat. # D-4522). Reactions were performed for 60 min at 30°C in FlashPlate Basic Microplate (PerkinElmer; Cat. # SMP 200) before getting stopped with 1% H_3_PO_4_. Plates were washed three times with 0.9% NaCl before [^33^P] incorporation was assessed by a scintillation counter. All steps involving [^33^P] were performed by ProQinase GmbH (Engesserstr. 4, 79108 Freiburg, Germany).

### Non- radioactive in vitro kinase assay

2 µg GST (GenScript; Cat. # Z02039-1), ubiquitin (R&D Systems; Cat. # U-100H), His-ubiquitin (R&D Systems; Cat. # U-530), HA-ubiquitin (R&D Systems; Cat. # U-110) or GST-ubiquitin (R&D Systems; Cat. # U-540) were incubated in 20 µl standard kinase reaction buffer (Cell Signaling Technology; Cat. # 9802S) containing 25 mM Tris-HCl (pH=7.5), 5 mM beta-glycerophosphate, 2 mM dithiothreitol (DTT), 0.1 mM Na_3_VO_4_, 10 mM MgCl_2_ which was supplemented with 1 mM ATP (Thermo Fisher; Cat. # A7699). PRKG1 assays additionally contain 1 μM cGMP (Merck; Cat. # G-6129). Final kinase concentration was 12.5 ng/µl for PRKACA (Thermo Fisher; Cat. # P2912), RPS6KA5 (Thermo Fisher; Cat. # PV3681), PRKG1 (Thermo Fisher; Cat. # PV4340), SGK2 (Thermo Fisher; Cat. # PV3858), FGFR1 (Promega; Cat. # V2991), FGFR1-V561M (Promega; Cat. # VA7456). His-MBP-PINK1 (R&D Systems, Cat. # AP-180-100) concentration was set to 50 ng/µl. Reactions were performed for 120 min at 30°C before reactions were stopped by Phos- tag™ sample buffer (FujiFilm Wako) and used for Phos-tag™ Coomassie and Phos-tag™ Immunoblot analysis.

### Coomassie

Proteins were supplemented with 4x NuPAGE™ sample buffer (Thermo Fisher; Cat # NP0008) and separated by NuPAGE™ 4–12% SDS-PAGE in precast gels (Thermo Fisher; Cat # NP0323BOX) with NuPAGE™ MES SDS Running Buffer (Thermo Fisher; Cat # NP0002). Gels were stained using Coomassie Brilliant Blue R-250 staining solution (BioRad; Cat # 1610437).

### Phos-tag™ Coomassie and Phos-tag™ Immunoblot

Proteins were supplemented with 3x Phos-tag™ sample buffer (FujiFilm Wako) and separated by 12.5% SuperSep™ Phos-tag™ precast gels (FujiFilm Wako; Cat # 199-18011) with running buffer (25 mM Tris, 192 mM glycine, 0.1% (wt/vol) SDS). For Coomassie: gels were stained using Coomassie Brilliant Blue R-250 staining solution (BioRad; Cat # 1610437). For Immunoblots: gels were soaked 30 min with ion removing buffer (25 mM Tris, 192 mM glycine, 1 mM EDTA, 10 % (vol/vol) methanol). Followed by 30 min incubation with transfer buffer (25 mM Tris, 192 mM glycine, 1mM EDTA, 10% (vol/vol) methanol). Proteins were transferred onto Immobilon-FL PVDF membranes (Millipore; Cat # IPFL00010), and nonspecific protein binding was blocked with 3% BSA in Tris-buffered saline for at least one hour at room temperature, followed by overnight incubation with specific primary antibodies in 3% BSA in Tris-buffered saline with 0.1% Tween-20 at 4°C. Primary antibodies were used as follows: GST (1:1,000 dilution; Cat # 27457701V) from Merck, ubiquitin (1:1,000 dilution; clone P4D1) from Santa Cruz Biotechnology. Membranes were washed three times with Tris-buffered saline containing 0.1% Tween-20 and incubated with the appropriate secondary antibodies coupled to IRDye 800CW/ IRDye 680RD (1:25,000 dilution; LI-COR Biosciences) or AzureSpectra- 490/AzureSpectra-550 (1:10,000 dilution; Azure Biosystems) for one hour at room temperature. Finally, membranes were washed three times with Tris-buffered saline containing 0.1 % Tween-20 and were analyzed with an Odyssey CLx imaging system (LI-COR Biosciences) or Sapphire Biomolecular Imager (Azure Biosystems).

### Mass spectrometry

Protein bands correlating to phosphorylated GST-ubiquitin were cut from SuperSep™ Phos- tag™ precast gels (FujiFilm Wako; Cat # 199-18011) and analyzed via mass spectrometry. In- gel trypsin digest was performed for all samples, and digested protein samples were measured on Q Exactive™ HF or Orbitrap Elite™. Data analysis was performed by Scaffold 4 (Proteome Software, Inc). Experiments were performed by the Core Facility for Mass Spectrometry & Proteomics (CFMP) at the Zentrum für Molekulare Biologie der Universität Heidelberg (ZMBH).

### AQUA mass spectrometry for tetra ubiquitin

AQUA mass spectrometry data for tetra ubiquitin was provided by R&D Systems. Lot # specific data can be obtained for K6 (Lot # 24369018A; Cat. # UC-15), K11 (Lot # 11068918A; Cat. # UC-45), K29 (Lot # 24571415; Cat. # UC-83), K33 (Lot # 02071217A; Cat. # UC-103), K48 (Lot # 33870318A; Cat. # UC-210B), K63 (Lot # 35670517B; Cat. # UC-310B) and pS65-ubiquitin (Lot # 22272117A; Cat. # UC-750).

### Phylogenetic kinase mapping

Phylogenetic tree for ubiquitin-specific kinases was created using Kinase Mapper (Reaction Biology).

### Protein alignment

Species-specific ubiquitin sequences were obtained from the following Uniprot-IDs: P0CG48; P0CG50; Q63429; P0CH28; P0CG69; P0CG63.

### In silico kinase prediction

Kinase/Phosphorylation site prediction was performed using NetPhos 3.1 Server (www.cbs.dtu.dk/services/NetPhos) using ensembles of neural networks.

### Databases

Tissue-specific gene expression and subcellular protein localization data for PRKCA, RPS6KA5, PRKG1 and PRKCB were obtained from Human Protein Atlas^31^.

## Extended Data Figures

**Extended Data Fig. 1:**
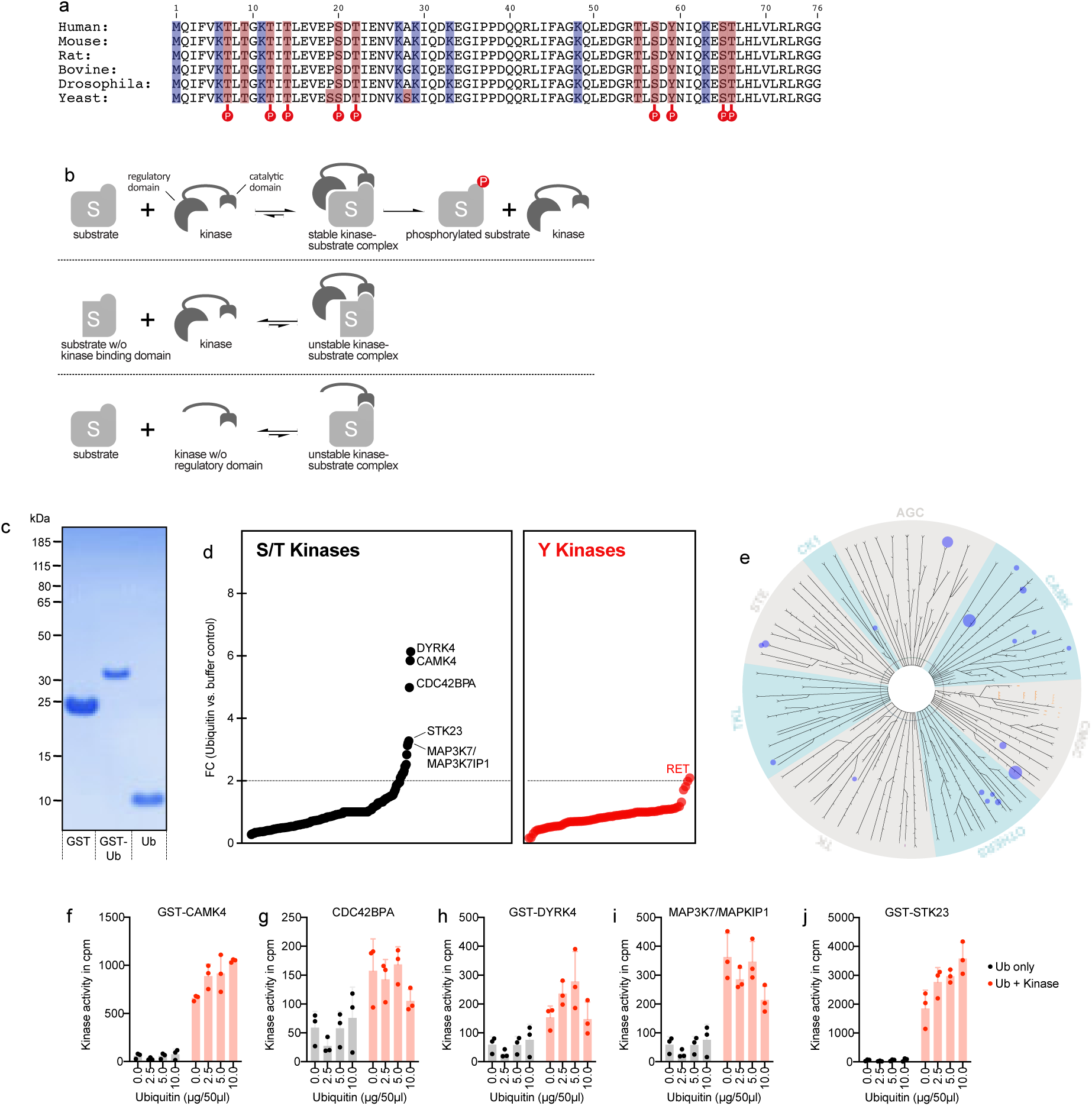
Kinase screening showed no phosphorylation for untagged ubiquitin. (a) Amino acid sequences of ubiquitin homologs. In blue amino acids responsible for additional ubiquitination are highlighted. Red indicates phosphorylatable amino acids. Phosphorylated sites are indicated based on data obtained from PhosphoSitePlus. (b) Schematic representation of kinase recruitment controlled by the regulatory kinase domain. The stabilization of the kinase-substrate complex by the regulatory kinase domain allows substrate phosphorylation. (c) SDS-PAGE of purified, recombinant GST, GST-ubiquitin (GST-Ub) and untagged ubiquitin (Ub). Proteins were visualized by Coomassie Brilliant Blue. (d) [^33^P] incorporation indicated as fold change (FC) between ubiquitin and buffer control. S/T kinases are shown in black and Y kinases are shown in red. Selected candidate kinases are named individually. Kinase screening was performed once. (e) Kinases with FC > 2, between untagged ubiquitin and protein-free control, are mapped on a phylogenetic kinase tree as blue dots. Dot size refers to FC measured in the kinase screening. (f-j) Candidate kinases were tested with increasing amounts of ubiquitin (0 - 10 µg/ 50 µL). Shown is the [^33^P] incorporation in counts per minute (cpm). Ubiquitin without kinase is shown in grey and ubiquitin with kinase in red. (individual data points are shown, bar graph: mean + SD, n= 3).

**Extended Data Fig. 2:**
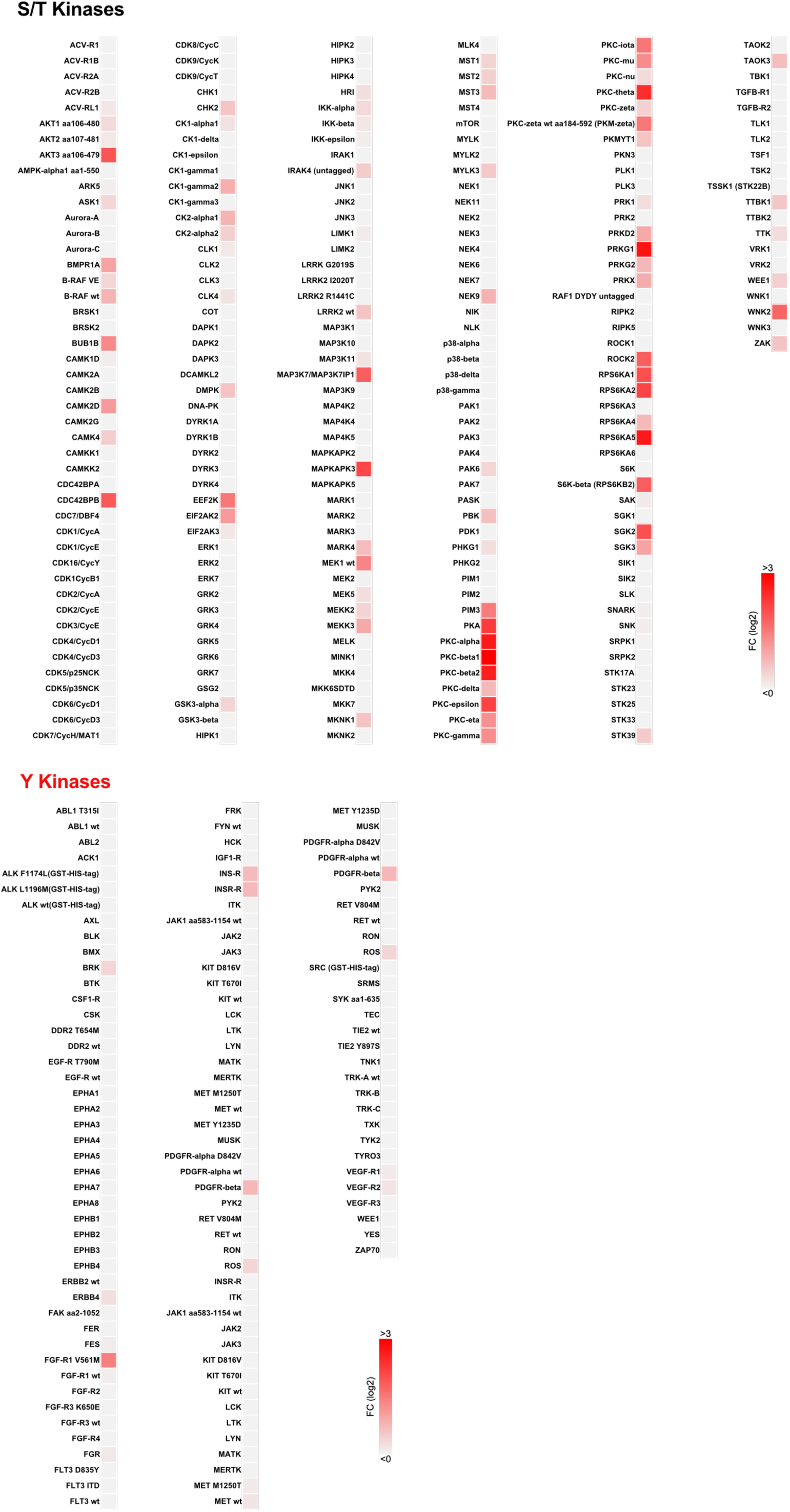
GST-ubiquitin phosphorylation by kinases. Alphabetical heat map indicating the ratio between [^33^P] incorporation of GST-ubiquitin and GST for each tested kinase. Upper panel lists S/T kinases, and the lower panel displays Y kinases. Kinase screening for GST and GST-ubiquitin was performed once.

**Extended Data Fig. 3:**
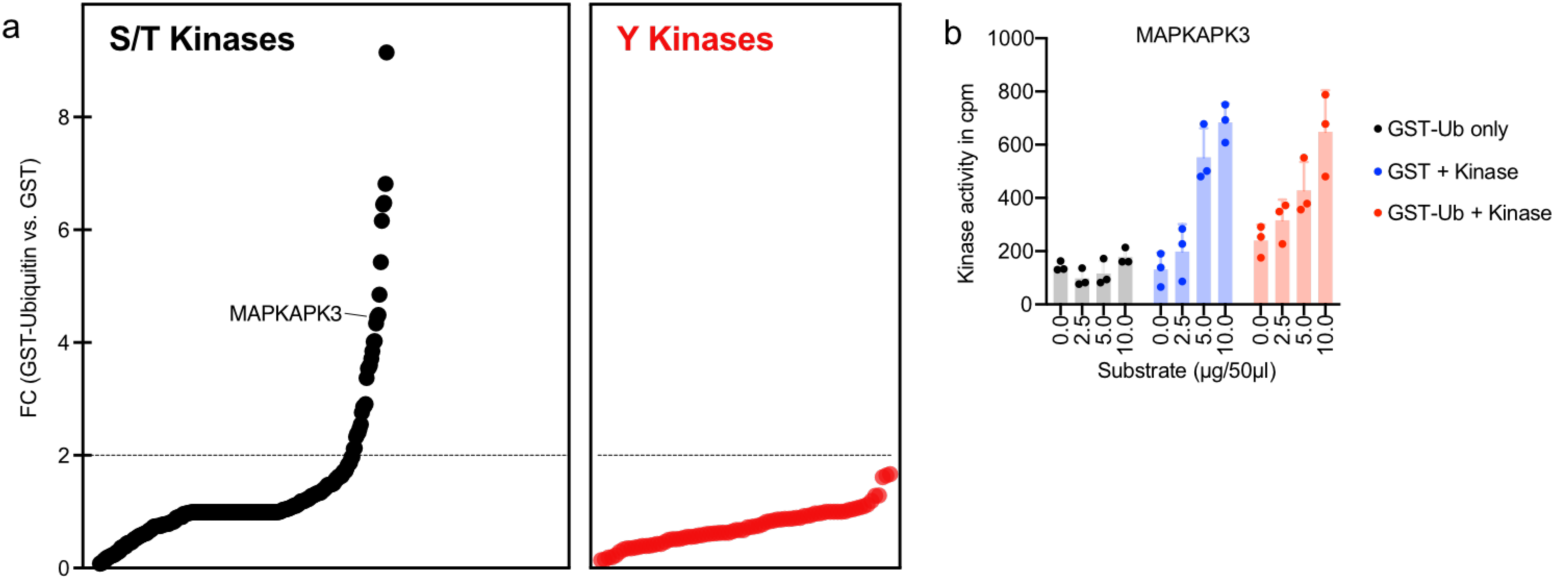
MAPKAPK3 does not phosphorylate GST-ubiquitin. (a) GST-ubiquitin phosphorylation measured by [^33^P] incorporation is displayed as FC between GST-ubiquitin and GST. MAPKAPK3 is indicated. Data is identical to [Fig. 1b] (b) Phosphorylation activity of MAPKAPK3 was tested with increasing amounts of GST-ubiquitin or GST (0 – 10 µg/ 50 µL). Shown is the [^33^P] incorporation in cpm. GST-ubiquitin without kinase is shown in grey, GST plus kinase in blue and conditions with GST-ubiquitin plus kinase in red. (individual data points are shown, bar graph: mean + SD, n= 3).

**Extended Data Fig. 4:**
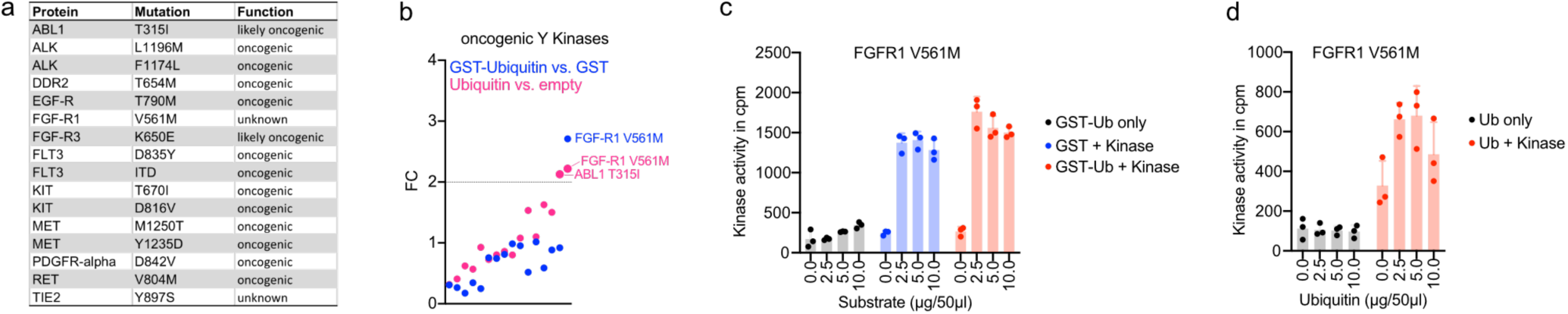
Oncogenic kinases show no signs of ubiquitin phosphorylation. (a) List of oncogenic kinase mutations tested in the ubiquitin phosphorylation screen. (b) [^33^P] incorporation indicated as fold change (FC) between ubiquitin and protein-free buffer (pink) or GST-ubiquitin and GST (blue) after incubation with oncogenic kinases listed in Extended Data Fig. 7a. Candidate oncogenic kinases with FC > 2 are named individually. Kinase screening was performed once. (c) Phosphorylation activity of FGFR1-V561M was tested with increasing amounts of GST-ubiquitin or GST (0 – 10 µg/ 50 µL). Shown is the [^33^P] incorporation in cpm. GST-ubiquitin without kinase is shown in grey, GST plus kinase in blue, GST-ubiquitin plus kinase in red. (individual data points are shown, bar graph: mean + SD, n= 3). (d) FGFR1-V561M substrate phosphorylation was tested with increasing amounts of ubiquitin (0 – 10 µg/ 50 µL). Shown is the [^33^P] incorporation in cpm. Ubiquitin without kinase is shown in grey and ubiquitin plus kinase in red. (individual data points are shown, bar graph: mean + SD, n= 3).

**Extended Data Fig. 5:**
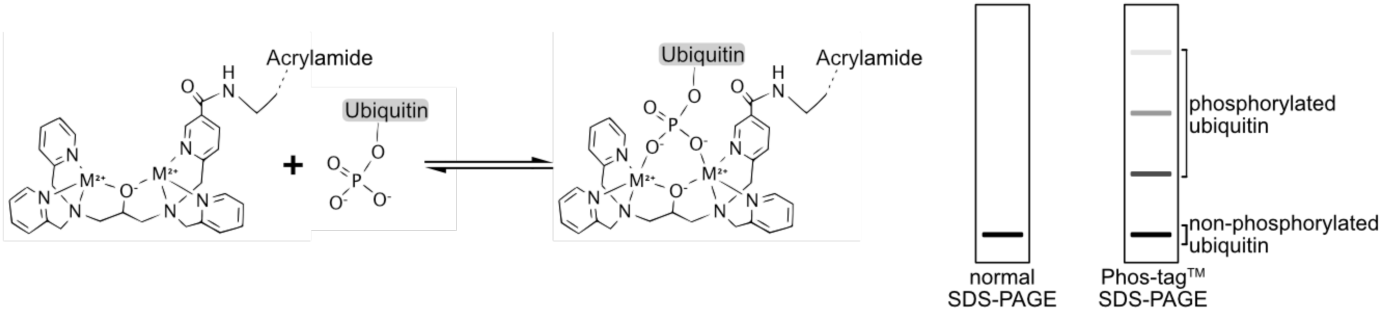
Phospho-ubiquitin detection by Phos-tag^™^ SDS-PAGE. Schematic representation of phospho-ubiquitin interaction with Phos-tag^™^ acrylamide. Interaction between phosphoryl group of phospho-ubiquitin and Phos-tag^™^ acrylamide reduces the migration speed during SDS-PAGE and creates additional protein bands which run higher compared to non-phosphorylated ubiquitin.

**Extended Data Fig. 6:**
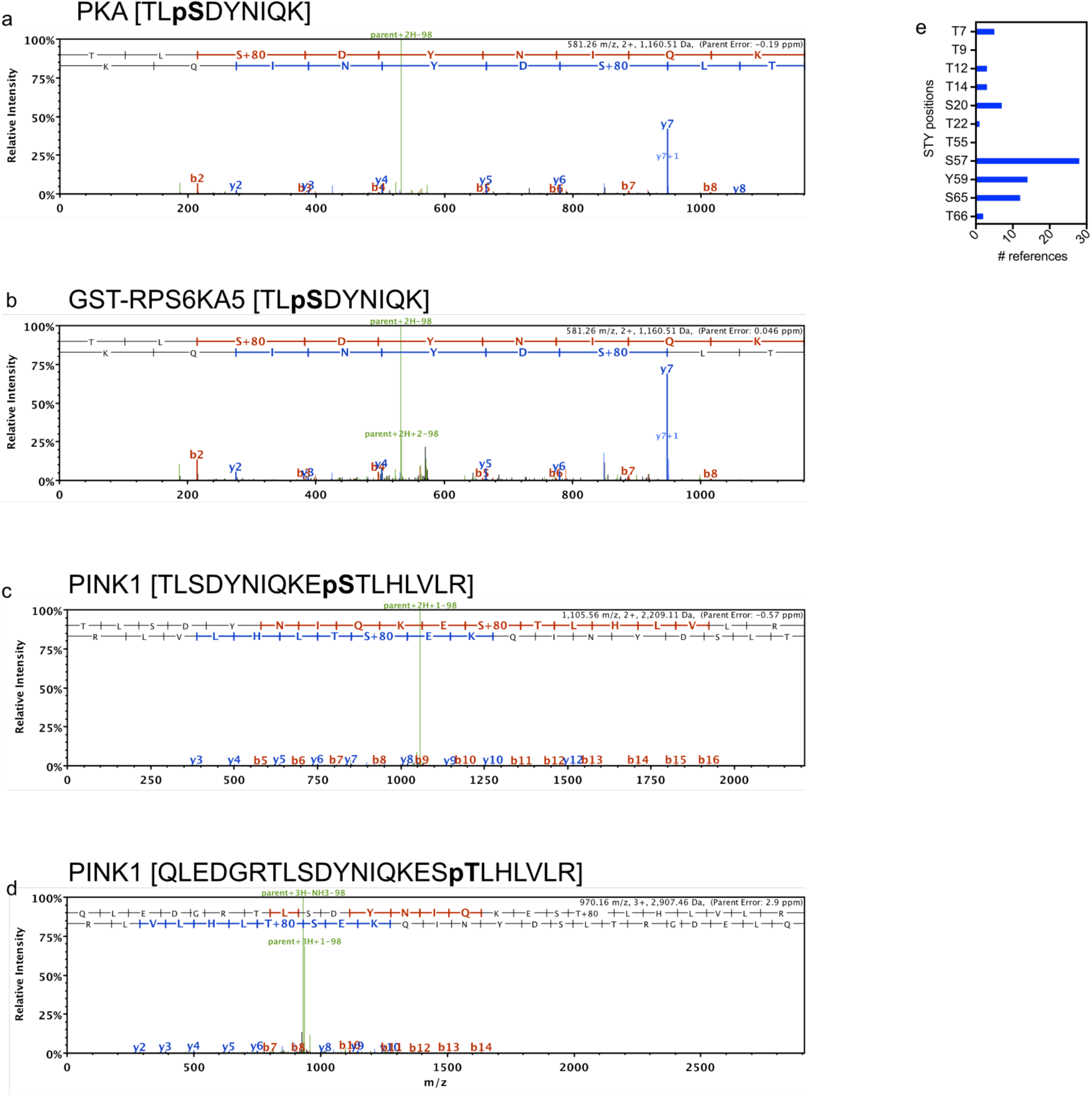
PKA and GST-RPS6KA5 phosphorylate ubiquitin at S57. (a-d) Mass spectrometry spectra, including predicted b- and y-ions for phosphorylated ubiquitin peptides. Kinases used for ubiquitin phosphorylation are indicated for each spectrum. (e) The Number of proteomic publications in which site-specific ubiquitin phosphorylations were observed. Data obtained from PhosphoSitePlus.

**Extended Data Fig. 7:**
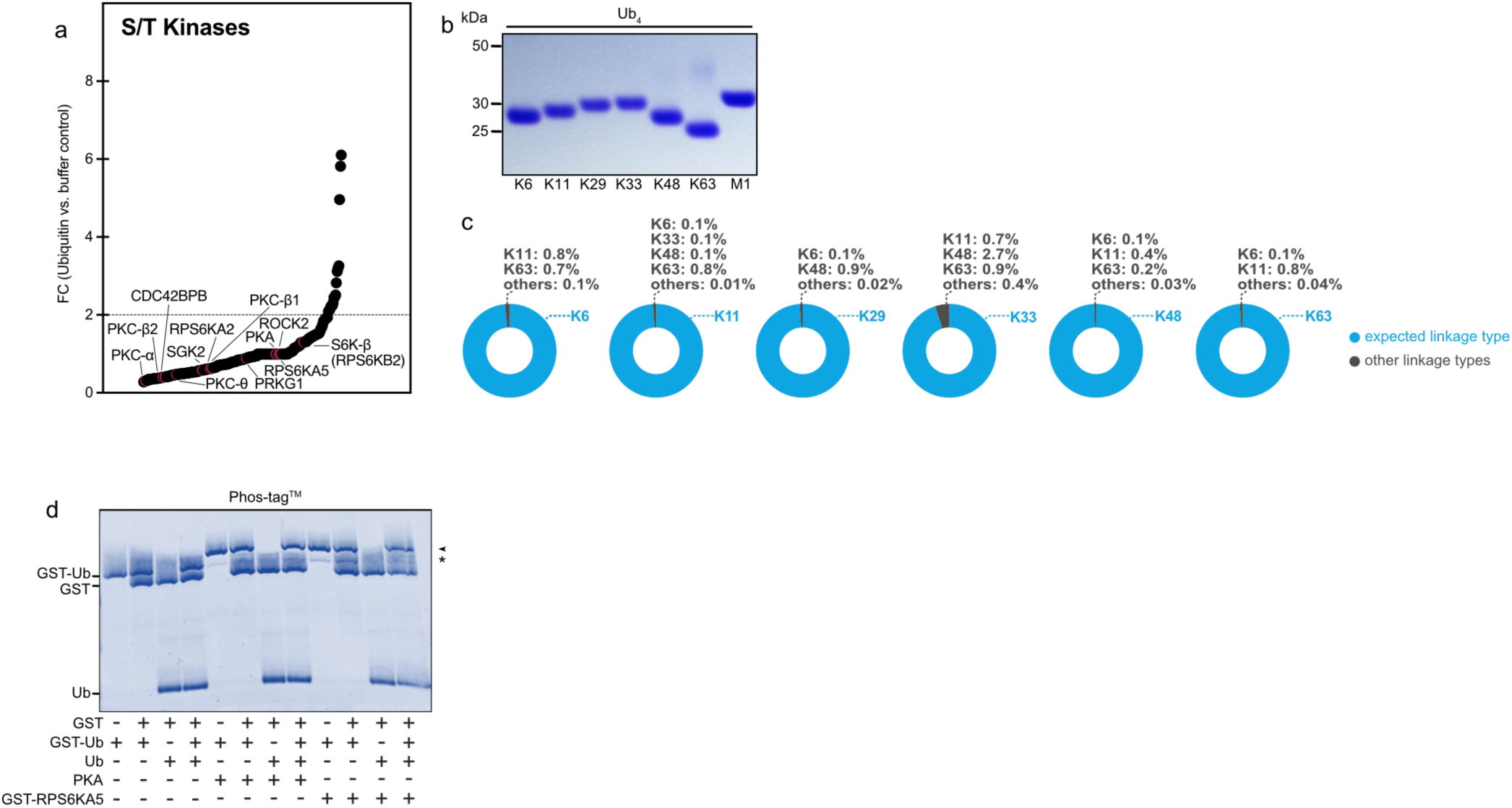
GST is not allosterically activating AGC kinases. (a) Ubiquitin phosphorylation displayed as [^33^P] incorporation FC between ubiquitin and protein-free control (data was previously shown in Extended Data Fig. 1d). Kinases phosphorylating GST-ubiquitin are indicated (compare Fig. 1b and Fig. 2). (b) Coomassie SDS- PAGE of tetra ubiquitin (Ub_4_) chains. (c) Donut chart representing the purity of Ub_4_ chains determined by AQUA mass-spectrometry. Intended ubiquitin linkage types are indicated in blue. Contaminating linkage types are shown in grey. (LOT specific data provided by Boston Biochem). (d) Coomassie analysis using Phos-tag^™^ SDS-PAGE to separate phosphorylated proteins after incubation with PKA or GST-RPS6KA5. GST, untagged ubiquitin and GST- ubiquitin were pooled as indicated, and PKA or GST-RPS6KA5 were added to the reactions. Arrow indicates phosphorylated GST-ubiquitin; Asterisk indicates non-phosphorylated GST- ubiquitin. Shown is a representative gel from three independent experiments.

**Extended Data Fig. 8:**
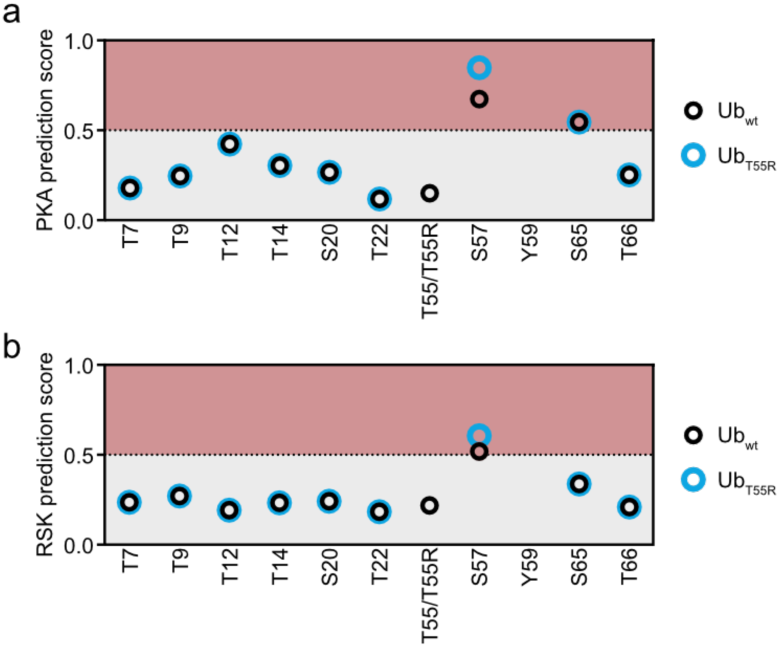
Amino acids surrounding S57 are suboptimal motifs for PKA and RSK. (a,b) PKA and RSK in silico phosphorylation prediction for each S/T within ubiquitin using NetPhos 3.1 server for Ub_wt_ and Ub_T55R_.

**Extended Data Fig. 9:**
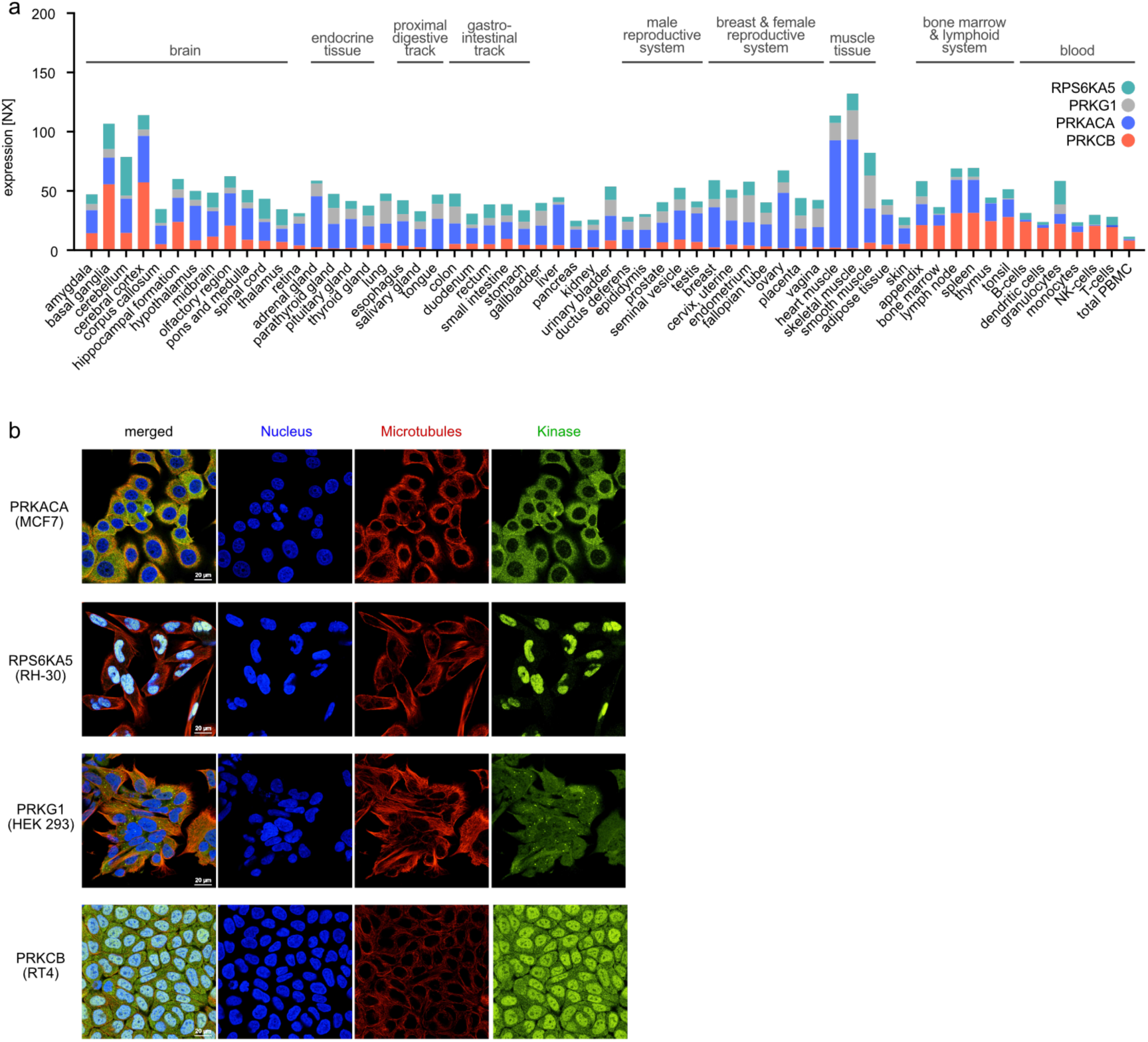
Universal expression of ubiquitin specific kinases. (a) Tissue-specific expression of PKA (PRKACA), RPS6KA5, PKG (PRKG1) and PKC-β1 (PRKCB) according to Human Protein Atlas. The expression is shown as consensus normalized expression (NX) levels. (b) Subcellular localization according to Human Protein Atlas for PKA (PRKACA), RPS6KA5, PKG (PRKG1) and PKC-β1 (PRKCB). Nuclei (blue), microtubules (red) and individual kinases (green) are shown. Visualized cell lines are indicated in brackets.

